# Integrating Machine-learning and Ultra-high-throughput Screening for Enzyme spaces exploration

**DOI:** 10.64898/2026.06.23.733994

**Authors:** Yitao Ke, Yanzhe Zhang, Minchao Fang, Jingyang Zhao, Hongli Zhu, Zehui Xu, Longxing Cao

## Abstract

The systematic navigation of biocatalyst space is constrained by elusive structure-activity rules and a lack of evolutionary history. Here, we present IMUSE, a strategy integrating machine learning with ultra-high-throughput screening. By screening millions of droplet-encapsulated *de novo* enzymes, we generated massive synthetic sequence-structure datasets to train models that capture their complex fitness landscapes and biophysical principles. These models effectively guide functional exploration across both sequence and novel structure spaces. IMUSE identified synergistic triple mutations yielding ∼5-fold activity improvements and discovered active second-generation designs with novel catalytic pockets, boosting the experimental success rate >4.9-fold (∼30%). This work demonstrates how synthetic fitness landscapes bridge the data gap in de novo enzyme space, transforming stochastic search into deterministic navigation to unlock highly proficient biocatalysts beyond natural boundaries.

## Introduction

Enzymatic catalysis is a kinetic process involving dynamic interaction between substrate and biocatalyst, yet evaluating and navigating the vast landscape of potential enzyme-substrate pairings with reactivity remains challenging (*1–4*). Currently, the development of reactivity progresses along two primary strategies (*5*). The first involves the remodeling of natural scaffolds via directed evolution or machine learning models, which have successfully leveraged evolutionary information to map sequence-activity relationships **(Fig. 1A)** (*6–8*). Nevertheless, these approaches are intrinsically confined to the local fitness landscapes of a limited number of ancestral folds, thereby restricting the structural diversity required to pair with the vast chemical space of unnatural substrates (*9*). The second strategy, *de novo* design, seeks to overcome these limitations by constructing bespoke scaffolds tailored to specific chemical targets (*10–13*). However, even with recent advances in artificial intelligence facilitating the generation of high-activity *de novo* enzymes inspired by native active sites, this vast, unexplored space lacks both evolutionary metadata and functional characterization, which thus impedes the establishment of effective strategies to systematically interrogate and navigate the functional scaffolds beyond known spaces (*14–17*).

**Figure 1.**
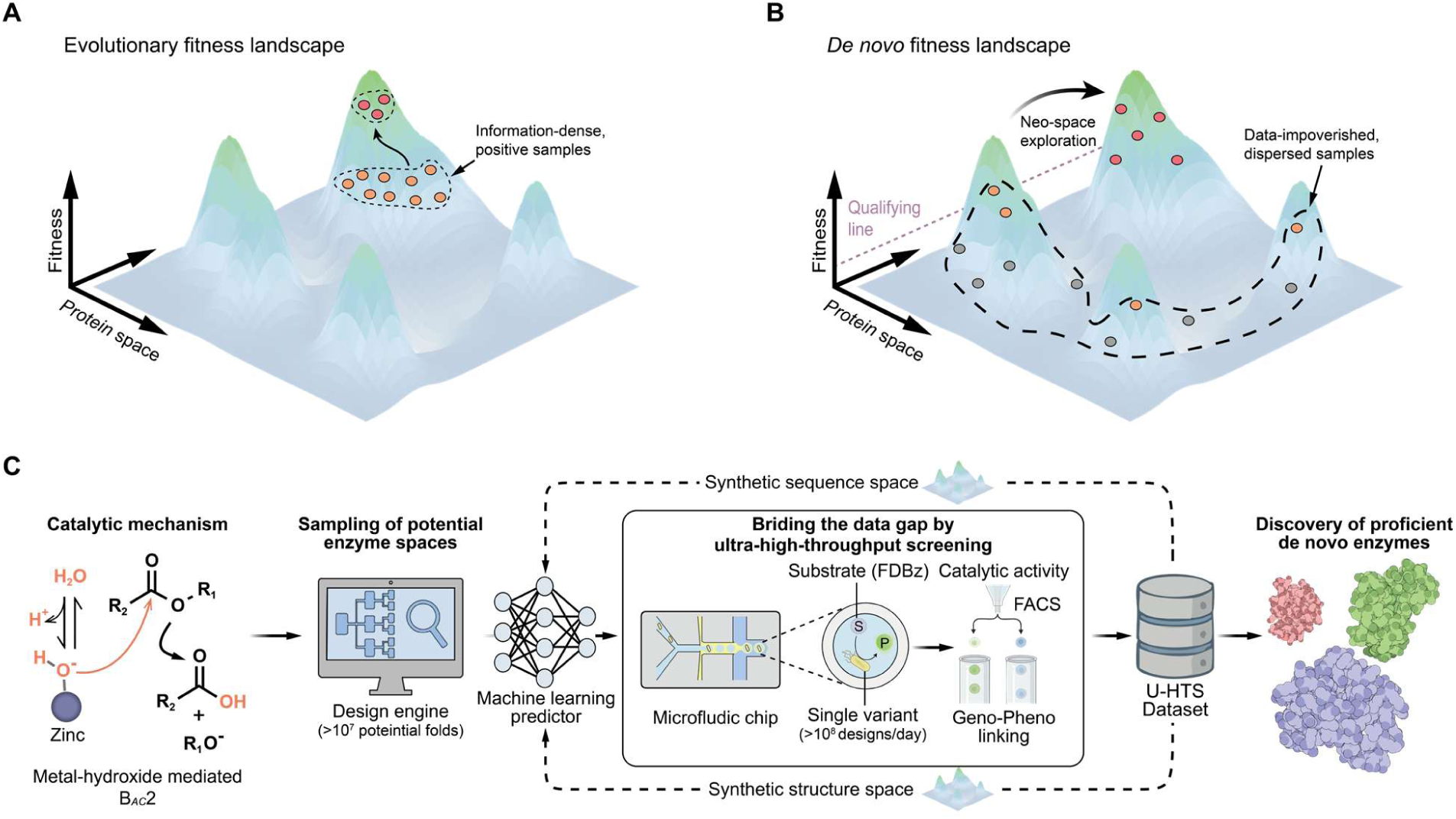
Present state of the method in biocatalyst exploration. (**A**) The evolutionary fitness landscape. Native enzyme families represent a compact but information-rich dataset which empowered the sequence space exploration. There is at present no method to navigate compatible enzymes beyond known superfamily, creating a huge hamper for their exploitation as key components in chemical process. (**B**) *De novo* enzymes are dispersed and remain a vast unexplored region of enzymes with uncharacterized biocatalytic activity, which inhibiting larger, non-intuitive leaps between fitness landscape. (**C**) IMUSE strategy for de novo biocatalyst spaces exploration. The Key steps in the IMUSE strategy. Step I: Scaffold sampling and sequence generation tailoring for diversifies chemo-physical favored theozymes. Step II: Designs are sorted with a sdDE-FACS platform that reflects catalytic activity. Active designs are isolated on the basis of fluorescent intensity on FACS. Step III: Machine learning models are trained on high-throughput dataset and applied to predict compatible enzymes.

Despite these challenges, the recent success of machine learning in connecting natural enzyme sequence space with diverse chemical substrates provides a compelling logic for exploring unknown biocatalyst structure space (*18*, *19*). However, connecting the small molecule to *de novo* enzymes is inherently more complex than mapping substrates to a given enzyme family, as the former requires navigating a high-dimensional landscape defined by vast topological breadth and immense sequence diversity (*20*, *9*). We reasoned that bridging the data gap would empower data-driven strategies to extend their success from native sequence space to the rational navigation of *de novo* sequence space, and even further into structure space **(Fig. 1B)** (*21*). Once a sufficient dataset was obtained, then machine learning models could be built to navigate between these two landscapes and enable the discovery of biocatalyst in a substrate-oriented fashion. However, since nature provides no “evolutionary map” for non-natural substrates, *de novo* spaces remain data-impoverished. Furthermore, these *de novo* enzymes feature stable folds and mechanistically reasonable active sites, which would make their activities indistinguishable by data-driven approaches utilizing native datasets. To overcome this bottleneck, we proposed the generation of “synthetic fitness landscapes” through ultra-high-throughput screening. This strategy emulates the data-dense environment of natural evolution by producing massive, substrate-specific functional datasets, effectively creating a roadmap for *de novo* enzyme space analogous to natural sequence databases.

In this study, we ask whether *de novo* enzyme space could be explored and exploited using machine learning. We developed a strategy called IMUSE (Integrating Machine-learning and Ultra-high-throughput Screening for Enzyme spaces exploration), which couples the sampling of low-energy scaffolds with ultra-high-throughput screening to generate data-dense functional repertoires spanning unexplored landscapes, thereby providing the empirical foundation for machine-learning model to identify functional patterns and navigate toward proficient *de novo* enzymes **(Fig. 1C)**. Isolating active designs within such massive libraries requires ultra-high-throughput screening methods, which can be achieved by generation of single droplet double emulsion (sdDE) (*22*). sdDE uses a microfluidic device to incorporate substrate and monoclonal variants that links genotype to catalytic phenotype at the single-cell level (*23*). To facilitate high-throughput screening, we selected zinc hydrolase as a model system and challenged it with synthetic fluorescein dibenzoate (FDBz) as the substrate. FDBz generates fluorescence upon hydrolysis, thereby serving as a quantitative readout of enzymatic activity. Ultimately, this platform yielded millions of droplets containing active *de novo* enzymes with diverse backbones, providing a foundational dataset to train a machine learning model for nominating active designs based on structural and sequence features. Applying the IMUSE, we bridged the data gap of *de novo* spaces and achieved substantial improvement in success rates in navigating both the sequence and structure landscapes, surpassing existing benchmarks. In sequence-space exploration, we identified variants whose catalytic efficiency was enhanced by approximately 5-fold through triple mutations that decouple kinetic constraints. Further, we constructed a second-generation library that samples a broader structural space, which yielded a higher proportion of active designs and whose characterized variants exhibited catalytic proficiency comparable to that of the native enzyme.

### Development of computational *de novo* enzyme design and ultra-high-throughput screening

To explore the vast and previously uncharted structural landscape, we developed a physics-based generative approach for *de novo* enzyme design, centered on the precise configuration of the theozyme for the target chemical reaction **(Fig. 2A)**. Specifically, for zinc-dependent catalysis, we systematically enumerated all possible histidine coordination geometries at the three available sites of the zinc ion, leaving one coordination site available to function as the general base for water activation—crucial for nucleophilic attack and ester bond cleavage (**fig. S1A, B**). For each coordinating histidine, all possible positions and rotameric conformations were generated and stored in a coordination RIF table (**fig. S1C**). To stabilize the transition state substrate, we further employed a RIF-based strategy to identify all favorable amino acid interactions with the substrate’s transition state geometry (**fig. S1D and table S2**). The spatial relationship between the transition state substrate and the coordinating histidines was systematically perturbed, allowing thorough sampling of theozyme configurations. Subsequently, a Monte Carlo fragment assembly approach was used to build protein backbones in a bottom-up manner, ensuring simultaneous accommodation of both the coordinating histidines and the transition state stabilizing residues (**fig. S1E**).

**Figure 2.**
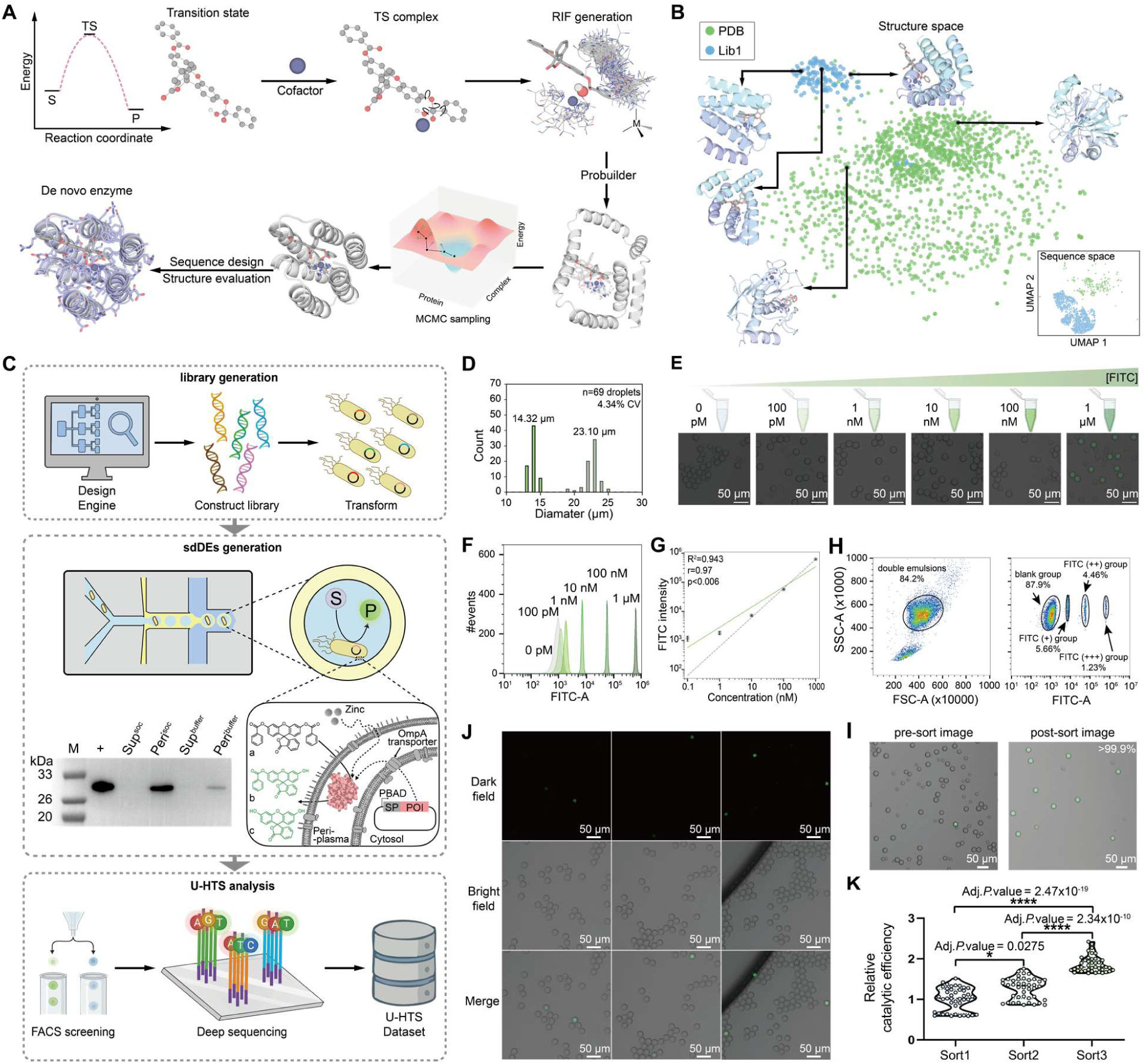
Ultra-high throughput single droplet isolation and proficient enzymes recovery. (**A**) *De novo* enzyme design starts from the substrate transition state calculated using quantum chemistry and compacted with perturbated physicochemical-satisfied coordination geometries generated with RIF generation (theozyme). ProBuilder samples protein fragments toward energetically improved conformations to support the theozyme, followed by sequence redesign with ProteinMPNN that encode the structure and stabilize the transition state. Designs are evaluated by structure prediction (for example, using AF2 and AF3) and are considered an in-silico success if the design (purple) and prediction (gray) align to a sufficient degree. (**B**) The method generates structurally diverse outputs visualized here by uniform manifold approximation and projection (UMAP). Bottom-right: sequence distribution between de novo zinc hydrolases from Lib1 and experimentally tested proteins sampled from PDB. Upper-left: structure distribution quantified here by qTM score between purified de novo zinc hydrolases from Lib1 (N = 145) and all clustered zinc protein families from PDB. There are only two representative Zn-His_3_ coordination folds in native protein. (**C**) Workflow of U-HTS dataset building. The fusion to the OmpA signal peptide enables efficient secretion of EGFP, which accumulates in the periplasmic protein fraction (‘Peri^soc^’) without contamination in the medium supernatant (‘Sup^soc^’). Prolonged incubation in buffer is confirmed no detectable leakage of secreted proteins (‘Peri^buffer^’ and ‘Sup^buffer^’). Purified EGFP (‘+’) was included as positive control. The marker (‘M’) of the first lane indicates the position of reference proteins of known mass. (**D**) Representative DE droplets size distributions (light green = inner diameter, dark green = outer diameter, CV = variation of total diameter). (**E**) Schematic and brightfield and fluorescence images of DE droplets containing multiple concentrations of hydrolyzed FDBz. (**F**) Histograms and relationship between measured intensities and concentration for DE droplets measured on the FACS. (**G**) The correlation in fluorescent intensity relative to multiple product concentrations measurements is plotted (R^2^ = 0.943, Pearson’s r = 0.97, P < 0.006). A gray dashed line represents the y = 0.001x line. The regression line is plotted as a solid green line. (**H-I**) FACS gates and post-sort image of droplets well with associated target enrichment sensitivity. (**J**) Prolonged incubation of enzymes within droplets. (**K**) Designs from original group and screened group were purified and assessed for catalytic efficiency, normalized by the average catalytic activity of original group. The screened designs had significantly better catalytic efficiency than the designs from the original group (\**P* < 0.05, \*\*\*\**P* < 0.0001; Kruskal-Wallis test with Dunn’s multiple comparisons). Data represent the mean ± s.d. of three independent measurements (G).

Structures featuring all three coordination histidines along with favorable transition state interactions were then selected for sequence optimization using Rosetta and ProteinMPNN (*24*, *25*). Top candidate designs were selected based on Rosetta energy metrics and further assessed for preorganization and structural stability using AlphaFold2 (AF2) (*26*). The resulting set of designs spans a broad range of protein architectures, encompassing a structurally diverse landscape. Evaluation with ESM2 and qTM scores against the Protein Data Bank, as well as Foldseek analysis, confirmed that our pipeline successfully samples enzymatic structural space beyond natural evolutionary boundaries, yielding a diverse array of *de novo* enzyme scaffolds (**Fig. 2B and fig. S2**) (*27*, *28*).

To overcome the bottleneck of functional verification for the generated massive *de novo* structures, we developed an ultra-high-throughput screening method by integrating single-droplet double emulsion with fluorescence-activated cell sorting (sdDE-FACS). This approach enabled the screening of millions of droplets within a single experimental run. Previous fluorescence-activated droplet sorting (FADS) methods have facilitated the screening of environmental metagenomes and engineered enzymes; however, while capable of assaying thousands of samples per second, their reliance on complex microfluidic setups remains a significant technical barrier to widespread application (*29–31*). We developed a custom microfluidic platform that generates FACS-compatible double emulsion (DE) droplets (**fig. S3**). Recognizing that prior DE systems often required additional lysis agents, which may complicate the discrimination of active enzymes, we constructed an arabinose-inducible periplasmic expression pathway in *E. coli* (*32*). This strategy compartmentalizes *de novo* enzymes within the periplasm to minimize interference from cytoplasmic proteins, thereby enabling accurate functional assessments **(Fig. 2C)**. To verify protein localization and retention within this compartment, we expressed EGFP in the periplasm and observed a good secretion of target without detectable leakage into the supernatant even with a prolonged incubation in reaction buffer **(Fig. 2C)**.

We designed a dual flow-focusing geometry chip system to generate highly monodisperse and stable DEs. In our system, the inner aqueous core co-encapsulates cells and substrates via an inlet tree with two widened channels, which minimizes cell-induced flow perturbations and enables immediate substrate-cell mixing, reducing the risk of product contamination across different designs (**fig. S1A**). This core is first emulsified in carrier oil to form water-in-oil (W/O) droplets, which are subsequently encapsulated within an outer aqueous phase to produce water-in-oil-in-water (W/O/W) DEs (**Fig. 2C** and **fig****. S3B)**. By systematically optimizing fluid flow rates, we generated uniformly DE droplets **(Fig. 2D, fig. S3D and fig. S4)**. Our chip-design ensures that the DE droplets remain significantly smaller than the FACS nozzle diameter to minimize deformation, while the high monodispersity effectively prevents stream instability and nozzle clogging, which are typically exacerbated by large variations in droplet size (*33*).

To ensure high assay specificity and sensitivity within these microcompartments, we set fluorescein dibenzoate (FDBz) as the fluorogenic substrate, as its low spontaneous hydrolysis minimizes background noise and allows for the precise discrimination of low-activity designs **(fig. S5)** (*34*). To quantify the dynamic range and lower limit of detection for sdDE-FACS, we loaded DE droplets with five concentrations of hydrolyzed FDBz and quantified emitted fluorescence via FACS on instruments. Microscopy images confirmed that DE droplets remained highly monodisperse when loaded with hydrolyzed FDBz **(Fig. 2E)**. Measured intensities for DE droplets cluster tightly as a function of loaded dye concentration, with both instruments clearly discriminating 10nM–1μM hydrolyzed FDBz from background and capable of detecting <1nM fluorescent probe (5 orders of magnitude) **(Fig. 2F, G)**. The compatibility of DE droplets with sdDE-FACS procedure was further validated using a 100 µm nozzle, where confocal microscopy confirmed that the DE droplets remained highly monodisperse and intact throughout the sorting process **(fig. 2H, I)**.

To validate the compatibility of active *de novo* enzymes with this sdDE-FACS procedure, we ultimately assessed the stability of cell encapsulated DE droplets by evaluating leakage and cross-talk during prolonged incubation. Confocal imaging revealed a distinct fluorescence contrast between positive and negative droplets with negligible fluorescence leakage, while enzymatic assays of enrichments further confirmed that the platform reliably distinguished active designs **(Fig. 2J, K)**. Collectively, these results establish sdDE-FACS as a foundational platform for the systematic exploration of the *de novo* landscape.

### High-throughput screening and structure validation of Lib1

With sdDE-FACS established, genes encoding the designed enzyme fragments were synthesized as oligo libraries and subsequently assembled into full-length genes using previously described method (*35*). The assembled library was then transformed into *E. coli* cells for functional screening at a throughput of thousands of droplets per second **(Fig. 2C)**. Individual *E. coli* cells expressing unique enzyme sequence were co-compartmentalized with zinc sulfate (ZnSO_4_) and fluorogenic substrate (FDBz) within microfluidic droplets. For the initial round, we applied permissive sorting gates to maximize the recovery of potential hits, followed sort rounds were performed under stringent conditions to enrich for highly active designs. The sorting pools were then subjected to deep-sequencing to identify potential hits. This workflow successfully isolated hundreds of active designs with diverse scaffolds from the vast background of non-functional sequences **(Fig. 3A)**. To assess whether existing computational metrics could navigate within data-impoverished spaces of these *de novo* enzymes, we evaluated the capability of physics-based metrics as well as deep learning-based scorers to distinguish between the active and inactive designs in Lib1 **(Fig. 3B and fig. S6)**. Among those computational scores, the most dominant physical feature is radius of gyration (Rg) and fa_atr_per_res_noLIG, which gauges protein compactness and correlates with stable packing. The other dominant features are AlphaFold3 based, relating to the compatibility of the protein and the substrate **(Fig. 3B)** (*36*). Collectively, deep-sequencing analysis of the sorted populations identified a weak enrichment of certain features, though it also indicates that the current scoring metrics are still unable to reliably distinguish active *de novo* enzymes (*37*).

**Figure 3.**
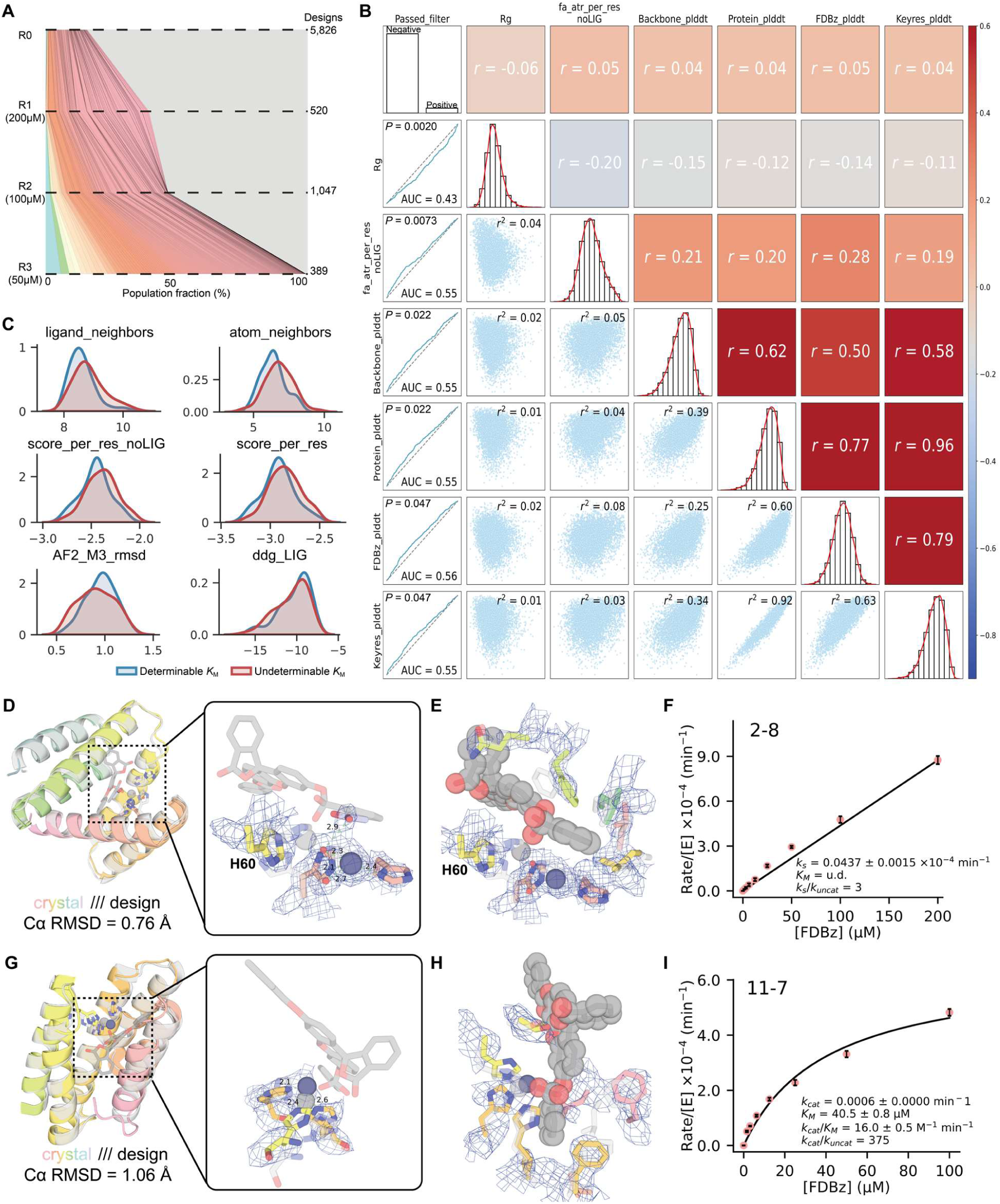
Generation and screening of de novo zinc hydrolase library, Lib1. (**A**) High-throughput evaluation of activity. Periplasmic-expressed active designs in *E. coli* are enriched during several rounds of selection at different substrate concentrations. The total number of unique designs identified in the NGS results after each round of sorting is indicated by the numbers on the right. The population fraction of each of the final surviving designs (with a population fraction larger than 0.0004 or enriched in the last round of selection) in each round of selection is shown as colored schemes. (**B**) Distribution of computational score between the final surviving designs (active) and discarded designs (inactive). These features show a statistically significance (FDR-adjusted *P* < 0.05, Mann-Whitney U test with Benjamini-Hochberg correction) between the final surviving designs (active) and discarded designs (inactive), but no feature is individually an effective discriminator. (**C**) Distribution of features between groups w/wo measurable *K*_M_. These features show different distributions between designs w/wo measurable *K*_M_, but no feature is individually an effective discriminator. Low values are favorable. (**D** and **G**) Structural superposition of design models (gray) and crystal structures (rainbow) for 2-8 (D) and 11-7 (G). Active site overlays of the design models (gray) and crystal structures (rainbow) of 2-8 (D) and 11-7 (G) with 2Fo − Fc map shown at 1.5σ and 1.0σ, respectively (blue mesh). (**E** and **H**) Superposition of substrate binding sites of the design models (gray) and crystal structures (rainbow) of 2-8 (E) and 11-7 (H) with 2Fo − Fc map shown at 1.5σ and 1.0σ, respectively (blue mesh). Distances shown in angstroms. (**F** and **I**) Michaelis-Menten plot for 2-8 (F) and 11-7 (I) with FDBz. Data represent the mean ± s.d. of three independent measurements of initial velocity (F, I).

To validate the screening results, we selected 145 unique sequences from the final enriched pool (343 in total) for individual purification and kinetic characterization **(fig. S7 to S10)**. All characterized monomeric designs displayed detectable hydrolytic activity. Kinetic characterization of these *de novo* enzymes exhibited rate accelerations (*k_cat_*/*k_uncat_*) ranging from 10^2^- to over 10^4^-fold relative to the uncatalyzed reaction (in the absence of enzyme) **(table S3)**. Among them, the top-performing variant, 3-4, displayed a catalytic proficiency of ∼1.6×10^8^ M^-1^ for FDBz, corresponding to a rate acceleration ∼ 10^4^-fold, which is comparable to a series of high active *de novo* designed metallohydrolases **(table S3)**.

However, while all designs were active, they exhibited varied kinetic behaviors. A subset of designs, typified by variant 2-8, showed clear catalytic turnover but failed to reach saturation kinetics within the solubility limit of the substrate, suggesting a defect in substrate affinity (*K_M_*) despite a functional catalytic center (**Fig. 3I**). Conversely, another group of designs displayed well-defined kinetic parameters for both substrate binding and turnover. To gain insight into the molecular features that impair the substrate affinity (*K_M_*), we compared score metrics w/o substrate affinity. We found, for example, that scores representing higher structural stability (score_per_res_w/oLIG), lower ligand neighbor atomic density (ligand_neighbors, atom_neighbors), as well as proper flexibility (AF2_M3_rmsd) and proper binding energy (ddg_LIG), were favored in many proficient (determinable *K_M_*) designs, which suggests balance of the global stability and pocket flexibility are required for enzyme design **(Fig. 3C and fig. S11)**. To understand the atomic-level basis of these distinct catalytic profiles, we solved the X-ray crystal structures of representative designs. The solved structures exhibited remarkable agreement with the computational models, with Cα root-mean-square deviations (RMSD) as low as 0.76 Å for design 2-8 and 1.06 Å for design 11-7 (calculated over 120 residues), thereby validating the high fidelity of the global backbone generated by our physicochemical-based generation method **(Fig. 3D, G and table S5)**.

The structure of 2-8 offered mechanistic insights into its decoupled affinity and catalysis. Within the active site of 2-8, an acetate molecule sits adjacent to the zinc ion, while a water molecule forms coordinating hydrogen bonds with the metal **(Fig. 3D)**. This specific arrangement remarkably mimics the nucleophilic attack on an ester bond, offering a structural basis for the preserved catalytic turnover observed in this variant. Further analysis revealed the structural origins of its poor substrate affinity **(Fig. 3E)**. The Zn (II)-coordinating histidine (H60) exhibited significant flexibility, wobbling away from the designed active site. In addition, rotamer shifts of other key histidine residues resulted in a steric clash with the coordinated zinc of the theozyme pose, which would thus lead to a steric clash between the substrate and the buried sidechains within this pocket. This lack of shape complementarity prevents effective substrate binding, consistent with the undetermined *K_M_* in kinetic assays **(Fig. 3F)**. In contrast, structural analysis of the high-affinity variant 11-7 revealed a precisely preorganized catalytic center even in the apo-state, which explains the measurable *K_M_* and high catalytic efficiency **(Fig. 3 G-I)**.

### Activity improvement within sequence spaces

Navigating across sequence landscapes, either naturally or through experimental approaches, has already proven effective in optimizing enzymes within evolutionary boundaries (*38*, *39*). However, the absence of evolutionary fitness information in *de novo* enzymes limits the efficacy of conventional optimization strategies, which typically rely on homologous sequence data to guide mutational trajectories (*14*, *40*). To explore the capability of IMUSE to navigate the sequence spaces of interest *de novo* enzymes, we selected two distinct active designs, 3-4 and 30-3, and performed random mutagenesis to systematically map their synthetic fitness landscapes **(Fig. 4B, E)**. We constructed libraries for both designs using error prone PCR and subjected them to ultra-high-throughput screening. This generated comprehensive functional maps for more than 10^5^ variants (ranging from single to triple mutations), providing compact yet information-rich datasets that reflect the functional preferences of the mutants and emulate the data-dense environment of natural evolution **(Fig. 4A, D and fig. S12)**. As the distal localization of most mutations suggests their effects may be mediated by stabilizing the active conformation or reshaping non-productive dynamics, detailed kinetic characterization further revealed that these designs were improved along distinct evolutionary trajectories to enhance catalytic performance (**fig. S13 to S16**) (*41*, *42*). For the 3-4 lineage, the library introduced mutations that significantly improved substrate affinity (*K_M_*), effectively locking the substrate into the pre-organized active site described above **(Fig. 4C and table S3)**. Comparatively, for the 30-3 lineage, optimization was driven by an enhancement in catalytic turnover number (*k_cat_*), suggesting the distinguished mutations that lowered the activation energy barrier or facilitated product release **(Fig. 4F and table S3)**. However, these distal mutations fail to decouple these parameters due to the lack of synergistic interactions required to break intrinsic constraints; in most cases, both *_kcat_*and *_KM_* increased concurrently, thereby negating any substantial gain in catalytic efficiency (*_kcat_/_KM_*), making activity improvements inefficient.

**Figure 4.**
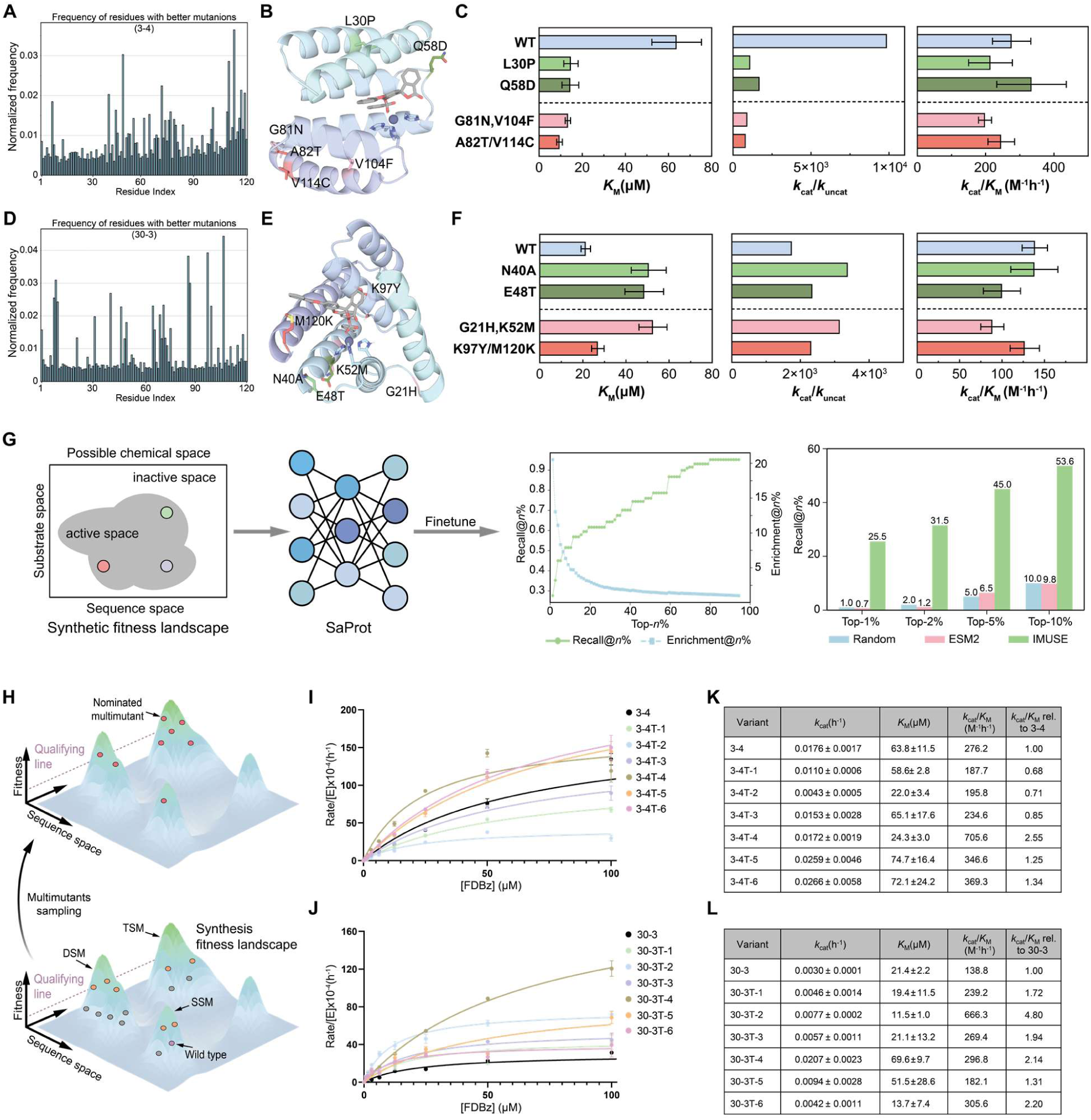
Improved performance within sequence space refinement. (**A** and **D**) Mutations identified from the 3-4 (A) and 30-3 (D) lineages by random mutations. (**B** and **E**) Design model of 3-4 (B) and 30-3 (E) in complex with FDBz substrate. Variants found in the site saturation mutation library are highlighted in the same color for each variant. (**C** and **F**) Comparisons of values of K_M_, k_cat_/k_uncat_ and k_cat_/K_M_ values for FDBz for four variants derived from 3-4 (C) and 30-3 (F), respectively. (**G**) A machine learning model was designed to map an enzyme sequence landscape of interest, identify high fitness mutants. Machine learning model based on IMUSE strategy were evaluated against the baseline model to calculate recall and enrichment at various degrees of *k*. (**H**) Schematic of our IMUSE strategy, that direct nominate hyperactive multimutants (red) within sequence space. Function-enhancing variants are identified with U-HTS screening, and after mapping fitness landscape through enriched (orange) and discarded (gray) mutants, a neural network is used to propose enhanced multimutants. SSM, single site mutation. DSM, double site mutation. TSM, triple site mutation. (**I, J**) Plot showing initial rates of enzyme-catalyzed hydrolysis of the FDBz substrate. (**K, L**) Kinetic constants for FDBz. Data represent the mean ± s.d. of three independent measurements of Michaelis–Menten parameters (C, F, K, L) and initial velocity (I, J).

To tackle the observed kinetic trade-offs, we developed a predictive machine learning framework capable of rationally navigating sequence space by integrating both sequence and structural features. Recognizing that sequence alone is insufficient to capture the specific mechanistic bottlenecks of diverse *de novo* scaffolds, we fine-tuned the structure-aware protein language model, SaProt, using our dense mutational datasets **(Fig. 4G and fig. S17A)** (*43*). In this pipeline, coarse structural tokens extracted from the representative backbones (3-4 and 30-3) via Foldseek were concatenated with mutation sequences to form comprehensive input representations (*28*). To counteract class imbalance within the high-throughput data and robustly distinguish highly functional variants, we employed a tailored batching strategy. By complementing the default cross-entropy loss with a supervised contrastive loss, the model effectively segregates the latent embeddings of distinct functional labels while clustering similar ones, thereby enhancing its resolution of the complex sequence-structure-function landscape.

We rigorously evaluated this strategy through retrospective benchmarking (80/20 training/test split), measuring the model’s capacity to generate a precisely ranked output list of candidate enzymes. Model performance was quantified using Precision@ *n*%, Enrichment@ *n*%, and Recall@*n*%, metrics that rigorously capture the fraction of true positive designs enriched at the very top of the prediction list. By effectively prioritizing favorable variants, our model enriched function-enhanced enzymes in the top 50 positions at a rate more than 50-fold higher than random sampling. Notably, this structure-informed strategy substantially outperformed predictions from the sequence-only ESM-2 model on this synthetic library (**Fig. 4G and fig. S17**) (*27*). Overall, our retrospective benchmarking validated the ability of IMUSE strategy to address the lack of evolutionary fitness information, providing an essential foundation for the effective sampling of higher-order multi-mutations.

With the ability to effectively nominate candidate multi-mutant variants, we next challenged IMUSE to sample the triple mutants (representing > 1.9×10^9^ theoretical combinations) and selected the top six variants for experimental validation (**Fig. 4H, fig. S18**

**to S20**). These top variants harbor mutations scattered across the protein structure, a pattern indicative of the long-range epistatic effects that drive natural enzyme evolution (*44*, *45*). Consistent with this synergistic mechanism, most of the nominated variants showed remarkable improvements in *_kcat_/_KM_*, achieving 2.6- to 4.8-fold over the wild-type enzyme and surpassing the previous best mutants by 2.1- to 4.8-fold **(Fig. 4I-L)**. This indicated that IMUSE successfully sampled mutations while preserving substantial gains in catalytic efficiency. Notably, IMUSE achieved higher sampling efficiency than a recent protein language model framework, MULTI-evolve, which requires 5- to 7- additive mutations to achieve comparable 1.5- to 5.8-fold improvement in *_k_*_cat_*/_K_*_M_ (*39*). Collectively, these findings demonstrate that our IMUSE strategy bridges the evolutionary data gap, enabling the rational mapping and navigation of *de novo* sequence landscapes to identify synergistic multi-mutation combinations that decouple kinetic constraints and enhance *de novo* enzyme performance.

### Navigating expanded structural space for active enzymes

Having established the capability for sequence space navigation, we proposed that this IMUSE could also be applied to navigate from a given structural space to an expanded space to distinguish potential active *de novo* enzymes. Previous retrospective analysis of Lib1 dataset revealed that none of the conventional metrics showed a significant correlation with experimental activity, highlighting the difficulty of evaluating *de novo* enzymes that lack evolutionary metadata (*37*). This failure of general scoring functions motivated us to develop a data-driven approach tailored to capture the complex fitness landscapes locally governing catalytic activity. We hypothesized that the functional data generated by IMUSE strategy could bridge this gap by learning features invisible to simple correlation models. To test this strategy, we adopted the broad genotype-phenotype data from the first-generation library (Lib1), establishing the first structural-level dataset of *de novo* enzyme–substrate complexes reported to date. Informed by the slight enrichment observed in AlphaFold3 confidence scores, we developed a geometric deep learning framework that integrates sequence embeddings from ESM2, residue-level pLDDT confidence scores from AlphaFold3, and structural representations extracted from design model (**Fig. 5A, fig. S21**) (*27*, *36*). Specifically, we used AlphaFold3 to predict and incorporate residue-level pLDDT scores as node-level features, enabling the model to explicitly account for structural confidence in AF3 predictions (*36*). These features, together with 3D atomic coordinates and residue identities, were represented as graphs in which spatial relationships were encoded using radial basis functions.

**Figure 5.**
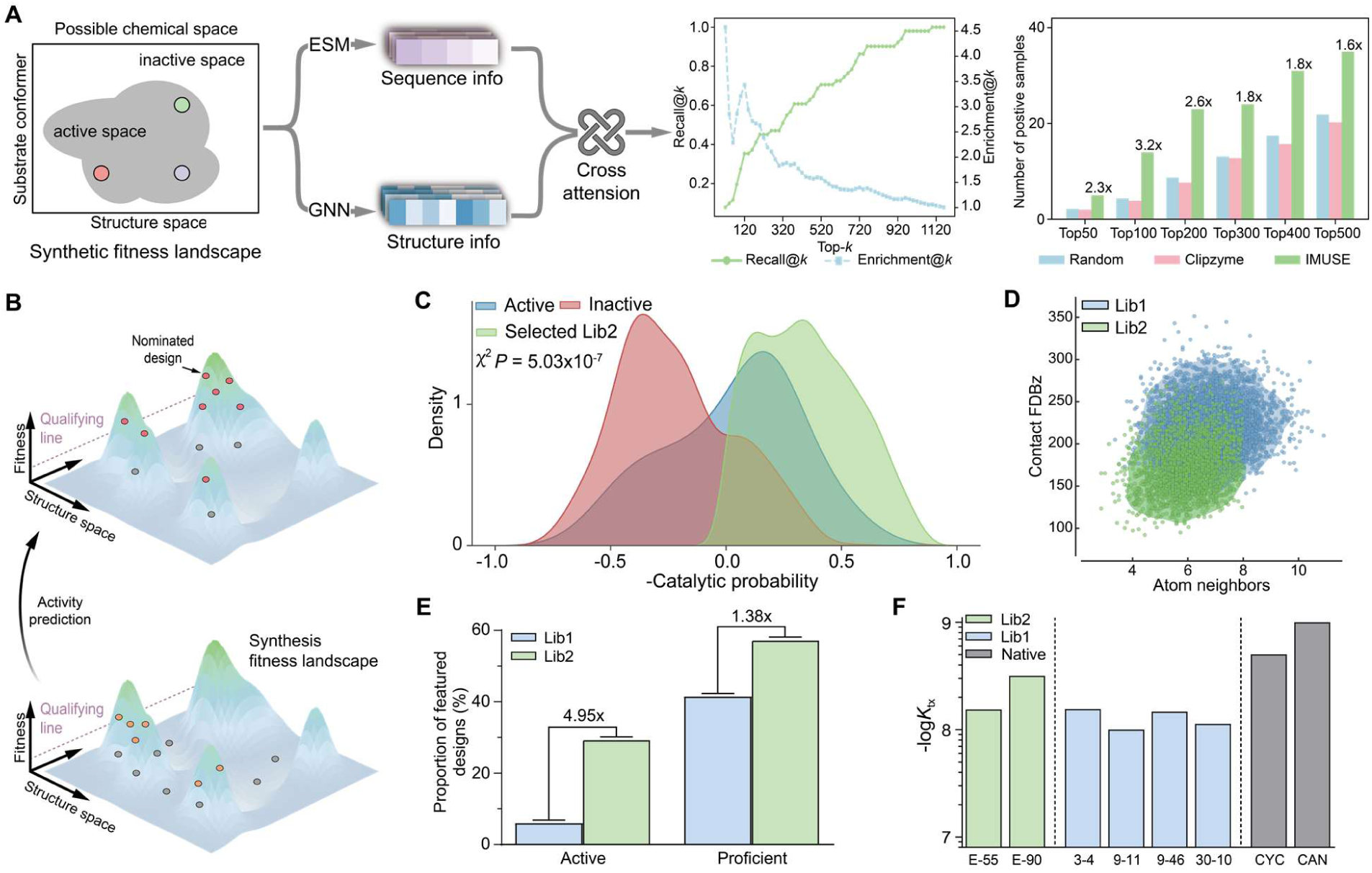
Building machine learning model for de novo structure spaces navigation in Lib2. (**A**) The process of building the machine learning model. The positive designs and negative designs from Lib1 were assessed with enrichments for the training dataset. Enzyme structures were encoded with pLDDT, sequences were processed with ESM2 model. Integration of structure and sequence were embedded by GNN network. Machine learning model based on IMUSE strategy were evaluated against state-of-the-art machine learning model by recall and enrichment at various degrees of k. (**B**) Schematic of IMUSE strategy, that nominate active designs (red) across structure space. Active designs are isolated with U-HTS screening, and after mapping fitness landscape through enriched (orange) and discarded (gray) designs, a neural network is applied for design activity prediction. (**C**) Separation of designs from test dataset recovered by sdDEs-FACS (blue) versus other designs (red) on the basis of the activity predictor (Mann-Whitney U test: P_(MWU)_ < 0.0001, Kolmogorov-Smirnov test: P_(KS)_ < 0.0001). In green, probability distribution for designs selected for the second-generation library. (**D**) Designs selected by the active predictor in Lib2 (green) and designs from Lib1 (blue). The physical features from these libraries exhibits marked differences. (**E**) Designs recovered in the second-generation library incorporate more active designs as well as proficient designs (with measurable *K*_M_) than in the first-generation library. (**F**) Catalytic proficiency of representative *de novo* zinc hydrolases from the Lib2 (green) and Lib1 (blue). The -log*K*_tx_ value of native enzymes extracted from ref (*47*) are plotted in gray. CYC, cyclophilin; CAN, carbonic anhydrase.

To integrate sequence and structural information, we fused sequence-level ESM2 embeddings with structural node features via a cross-attention mechanism (*27*). Notably, for zinc-coordinated scaffolds, we introduced a dedicated attention module that prioritizes the catalytic environment, allowing the model to capture local coordination geometry and functionally dominant interactions. The overall architecture employs a GVP-GNN encoder to perform geometric message passing and generate graph-level embeddings for activity prediction. Finally, a classification head was used to predict binary outputs (True/False), which were further used to derive a ranking score.

To validate the activity predictor, we applied benchmarking experiments to evaluate its overall effectiveness in enrichment of active enzymes from the initial library. In consideration of the low ratio of recovered designs in the Lib1, we implemented an 80/20 training/test split for this retrospectively analysis. This model was able to enrich the predictions ranked in the top fifty positions with enzymes with catalytic activity >2-fold better than would be observed by randomly sampling Lib1, and also surpassed baseline enzyme prediction methods (Clipzyme) (*46*). To gain insight into the molecular features of the active and inactive designs identified by our model, comparative analysis further revealed significant structural divergences, suggesting that the model implicitly converged on fundamental biophysical principles rather than simply memorizing training examples; it learned to penalize structural feratures associated with unfavored catalytic process, observed as a better performance on prediction of complex model **(fig. S22)**. To further improve model performance, we next performed ablation study on the test dataset to identify key features that contribute to this activity predictor **(fig. S23)**.

Encouraged by this success in the initial library, we next asked whether the model built from the first-generation library could generalize to navigate entirely unknown structural spaces (**Fig. 5B**). To rigorously test this, we updated the activity predictor with full dataset from Lib1. Subsequently, we generated a second-generation library (Lib2) using the same physics-based generative approach to ensure structural diversity. We then ranked the designs based on the computational metrics as Lib1 and selected them according to our activity predictor combined with key features identified in Lib1 (**Fig. 5C**). This procedure nominated thousands of designs with expanded structural diversity, incorporating diverse catalytic pocket and substrate interaction beyond the first library space **(Fig. 5D)**. The generalizability of IMUSE strategy was ultimately confirmed by the increased success rate, as we synthesized and purified 96 candidates from Lib2 to identify 28 active designs (29.2%), with ∼4.9-fold increase in the rate of active hits compared to that of the previous Lib1 **(Fig. 5E)**. These results underscore that the model built on high-throughput screening data can be applied to effectively select a diverse and active set of designs, enabling effective navigation within an expanded *de novo* design landscape.

Next, we focused on these active designs for detailed characterization (**fig. S24 to S27**). Strikingly, 16 of these 28 designs (∼57%) exhibited measurable binding affinity and catalytic activity toward the substrate **(Fig. 5E, fig. S26)**, which stands in contrast to the 40% success rate typically observed in Lib1. Kinetic characterization further revealed that two of these model-selected designs displayed catalytic proficiency (-log *K_tx_*) exceeding the most active designs from the first library and even comparable to native enzyme **(Fig. 5F, fig. S28 and table S3)** (*47*). These results represent a marked success in the exploration of data-impoverished structural space for *de novo* enzyme design, and they underscore that the lessons we have learned from high-throughput screening can be applied to generate a diverse and highly active set of designs suitable for either high- or low-throughput screening.

## Discussion

Systematic exploration of the *de novo* structural landscape is a prerequisite for advancing biocatalysis, yet the absence of evolutionary metadata and landscape-activity complexity have historically limited the navigation of these vast spaces (*21*, *37*, *38*, *48*). IMUSE firstly address this conflict by effectively emulating the data-dense environment of natural evolution, generating the synthetic fitness landscapes required to bridge the gap between computational design and functional realization. Our approach provides a structural diversity that significantly exceeds the limitations of natural metagenomic libraries, which enables the interrogation of entirely novel landscapes, transcending the phylogenetic constraints of traditional enzyme engineering. Current methods for developing biocatalysts for unnatural substrate rely on remodeling from natural scaffolds or screening from *de novo* designs (*17*). Constrained by number of native folds and screening efficiency, these methods validate a small fraction of sequence and structure space, whereas we show in this system that IMUSE can not only identified variants with order-of-magnitude increases in catalytic efficiency, but were also capable of exploring millions of structurally diverse designs of which over thousands of them are recovered with catalytic activity.

Furthermore, we demonstrate that sdDE-FACS is an effective strategy for the high-throughput isolation of active *de novo* enzymes by coupling genotype to phenotype, and that it could be extended to either natural or engineered enzymes catalyzing other reaction types (*49–51*). The combination of *de novo* enzyme design and sdDE-FACS enabled the implementation of effective design-test-learn cycles, yielding a deeper understanding of the design principles for *de novo* enzymes as previously validated in *de novo*–designed miniproteins (*52*, *53*). This platform provided a foundational framework to accelerate the discovery of *de novo* biocatalysts across previously inaccessible regions of the protein universe and would advance the development of next-generation, AI-based enzyme design methods.

## Notes

### Competing Interest Statement

The authors have declared no competing interest.

## References

1. S. J. Benkovic, S. Hammes-Schiffer, A Perspective on Enzyme Catalysis. Science 301, 1196–1202 (2003).

2. G. P. Pinto, M. Corbella, A. O. Demkiv, S. C. L. Kamerlin, Exploiting enzyme evolution for computational protein design. Trends in Biochemical Sciences 47, 375–389 (2022).

3. J. Yang, F.-Z. Li, F. H. Arnold, Opportunities and Challenges for Machine Learning-Assisted Enzyme Engineering. ACS Cent. Sci. 10, 226–241 (2024).

4. J. Listgarten, H. Jiang, How artificial intelligence is reengineering protein engineering. Science 392, 159–166 (2026).

5. E. L. Bell, A. E. Hutton, A. J. Burke, A. O’Connell, A. Barry, E. O’Reilly, A. P. Green, Strategies for designing biocatalysts with new functions. Chem. Soc. Rev. 53, 2851–2862 (2024).

6. K. K. Yang, Z. Wu, F. H. Arnold, Machine-learning-guided directed evolution for protein engineering. Nat Methods 16, 687–694 (2019).

7. N. S. Sarai, T. J. Fulton, R. L. O’Meara, K. E. Johnston, S. Brinkmann-Chen, R. R. Maar, R. E. Tecklenburg, J. M. Roberts, J. C. T. Reddel, D. E. Katsoulis, F. H. Arnold, Directed evolution of enzymatic silicon-carbon bond cleavage in siloxanes. Science 383, 438–443 (2024).

8. H. Seo, H. Hong, J. Park, S. H. Lee, D. Ki, A. Ryu, H.-Y. Sagong, K.-J. Kim, Landscape profiling of PET depolymerases using a natural sequence cluster framework. Science 387, eadp5637 (2025).

9. L. H. Isakova, E. Streltsova, O. O. Bochkareva, P. K. Vlasov, F. A. Kondrashov, Descent from a common ancestor restricts exploration of protein sequence space. Proceedings of the National Academy of Sciences 123, e2532018123 (2026).

10. A. H.-W. Yeh, C. Norn, Y. Kipnis, D. Tischer, S. J. Pellock, D. Evans, P. Ma, G. R. Lee, J. Z. Zhang, I. Anishchenko, B. Coventry, L. Cao, J. Dauparas, S. Halabiya, M. DeWitt, L. Carter, K. N. Houk, D. Baker, De novo design of luciferases using deep learning. Nature 614, 774–780 (2023).

11. A. Lauko, S. J. Pellock, K. H. Sumida, I. Anishchenko, D. Juergens, W. Ahern, J. Jeung, A. F. Shida, A. Hunt, I. Kalvet, C. Norn, I. R. Humphreys, C. Jamieson, R. Krishna, Y. Kipnis, A. Kang, E. Brackenbrough, A. K. Bera, B. Sankaran, K. N. Houk, D. Baker, Computational design of serine hydrolases. Science 388, eadu2454 (2025).

12. W. Ahern, J. Yim, D. Tischer, S. Salike, S. M. Woodbury, D. Kim, I. Kalvet, Y. Kipnis, B. Coventry, H. R. Altae-Tran, M. S. Bauer, R. Barzilay, T. S. Jaakkola, R. Krishna, D. Baker, Atom-level enzyme active site scaffolding using RFdiffusion2. Nat Methods 23, 96–105 (2026).

13. M. Braun, A. Tripp, M. Chakatok, S. Kaltenbrunner, C. Fischer, D. Stoll, A. Bijelic, W. Elaily, M. G. Totaro, M. Moser, S. Y. Hoch, H. Lechner, F. Rossi, M. Aleotti, M. Hall, G. Oberdorfer, Computational enzyme design by catalytic motif scaffolding. Nature 649, 237–245 (2026).

14. F. H. Arnold, Directed Evolution: Bringing New Chemistry to Life. Angewandte Chemie International Edition 57, 4143–4148 (2018).

15. J. A. Vila, About the Protein Space Vastness. Protein J 39, 472–475 (2020).

16. C. Jurich, Q. Shao, X. Ran, Z. J. Yang, Physics-based modeling in the new era of enzyme engineering. Nat Comput Sci 5, 279–291 (2025).

17. C. W. Kosonocky, S. Alamdari, K. K. Yang, A. P. Amini, Closing the loop: Experimentally validated methods in artificial intelligence–driven protein design. Current Opinion in Structural Biology 98, 103272 (2026).

18. A. E. Paton, D. A. Boiko, J. C. Perkins, N. I. Cemalovic, T. Reschützegger, G. Gomes, A. R. H. Narayan, Connecting chemical and protein sequence space to predict biocatalytic reactions. Nature 646, 108–116 (2025).

19. H. Cui, Y. Su, T. J. Dean, T. Yu, Z. Zhang, J. Peng, D. Shukla, H. Zhao, Enzyme specificity prediction using cross-attention graph neural networks. Nature 647, 639–647 (2025).

20. I. S. Povolotskaya, F. A. Kondrashov, Sequence space and the ongoing expansion of the protein universe. Nature 465, 922–926 (2010).

21. K. K. Yang, Z. Wu, F. H. Arnold, Machine-learning-guided directed evolution for protein engineering. Nat Methods 16, 687–694 (2019).

22. R. Buller, S. Lutz, R. J. Kazlauskas, R. Snajdrova, J. C. Moore, U. T. Bornscheuer, From nature to industry: Harnessing enzymes for biocatalysis. Science 382, eadh8615 (2023).

23. M. Gantz, S. Neun, E. J. Medcalf, L. D. van Vliet, F. Hollfelder, Ultrahigh-Throughput Enzyme Engineering and Discovery in In Vitro Compartments. *Chem. Rev*. 123, 5571–5611 (2023).

24. M. D. Tyka, D. A. Keedy, I. André, F. DiMaio, Y. Song, D. C. Richardson, J. S. Richardson, D. Baker, Alternate States of Proteins Revealed by Detailed Energy Landscape Mapping. Journal of Molecular Biology 405, 607–618 (2011).

25. J. Dauparas, I. Anishchenko, N. Bennett, H. Bai, R. J. Ragotte, L. F. Milles, B. I. M. Wicky, A. Courbet, R. J. de Haas, N. Bethel, P. J. Y. Leung, T. F. Huddy, S. Pellock, D. Tischer, F. Chan, B. Koepnick, H. Nguyen, A. Kang, B. Sankaran, A. K. Bera, N. P. King, D. Baker, Robust deep learning–based protein sequence design using ProteinMPNN. Science 378, 49–56 (2022).

26. J. Jumper, R. Evans, A. Pritzel, T. Green, M. Figurnov, O. Ronneberger, K. Tunyasuvunakool, R. Bates, A. Žídek, A. Potapenko, A. Bridgland, C. Meyer, S. A. A. Kohl, A. J. Ballard, A. Cowie, B. Romera-Paredes, S. Nikolov, R. Jain, J. Adler, T. Back, S. Petersen, D. Reiman, E. Clancy, M. Zielinski, M. Steinegger, M. Pacholska, T. Berghammer, S. Bodenstein, D. Silver, O. Vinyals, A. W. Senior, K. Kavukcuoglu, P. Kohli, D. Hassabis, Highly accurate protein structure prediction with AlphaFold. Nature 596, 583–589 (2021).

27. Z. Lin, H. Akin, R. Rao, B. Hie, Z. Zhu, W. Lu, N. Smetanin, R. Verkuil, O. Kabeli, Y. Shmueli, A. dos Santos Costa, M. Fazel-Zarandi, T. Sercu, S. Candido, A. Rives, Evolutionary-scale prediction of atomic-level protein structure with a language model. Science 379, 1123–1130 (2023).

28. M. van Kempen, S. S. Kim, C. Tumescheit, M. Mirdita, J. Lee, C. L. M. Gilchrist, J. Söding, M. Steinegger, Fast and accurate protein structure search with Foldseek. Nat Biotechnol 42, 243–246 (2024).

29. P. Gruner, Y. Skhiri, B. Semin, Q. Brosseau, A. D. Griffiths, V. Taly, J.-C. Baret, Microfluidic Approaches for the Study of Emulsions: Transport of Solutes. MRS Online Proceedings Library 1530, 201 (2013).

30. A. Sciambi, A. R. Abate, Accurate microfluidic sorting of droplets at 30 kHz. Lab Chip 15, 47–51 (2014).

31. R. H. Cole, S.-Y. Tang, C. A. Siltanen, P. Shahi, J. Q. Zhang, S. Poust, Z. J. Gartner, A. R. Abate, Printed droplet microfluidics for on demand dispensing of picoliter droplets and cells. Proceedings of the National Academy of Sciences 114, 8728–8733 (2017).

32. M. Jeschek, R. Reuter, T. Heinisch, C. Trindler, J. Klehr, S. Panke, T. R. Ward, Directed evolution of artificial metalloenzymes for in vivo metathesis. Nature 537, 661–665 (2016).

33. S. Ma, J. M. Sherwood, W. T. S. Huck, S. Balabani, The microenvironment of double emulsions in rectangular microchannels. Lab Chip 15, 2327–2334 (2015).

34. H. Fengying, W. Liufang, L. Bingrui, W. Qiguang, W. Qi, Studies on crystal structure and hydrolysis features of the fluorescein dibenzoate. Acta Chimica Sinica 51, 119 (1993).

35. J. C. Klein, M. J. Lajoie, J. J. Schwartz, E.-M. Strauch, J. Nelson, D. Baker, J. Shendure, Multiplex pairwise assembly of array-derived DNA oligonucleotides. Nucleic Acids Res 44, e43 (2016).

36. J. Abramson, J. Adler, J. Dunger, R. Evans, T. Green, A. Pritzel, O. Ronneberger, L. Willmore, A. J. Ballard, J. Bambrick, S. W. Bodenstein, D. A. Evans, C.-C. Hung, M. O’Neill, D. Reiman, K. Tunyasuvunakool, Z. Wu, A. Žemgulytė, E. Arvaniti, C. Beattie, O. Bertolli, A. Bridgland, A. Cherepanov, M. Congreve, A. I. Cowen-Rivers, A. Cowie, M. Figurnov, F. B. Fuchs, H. Gladman, R. Jain, Y. A. Khan, C. M. R. Low, K. Perlin, A. Potapenko, P. Savy, S. Singh, A. Stecula, A. Thillaisundaram, C. Tong, S. Yakneen, E. D. Zhong, M. Zielinski, A. Žídek, V. Bapst, P. Kohli, M. Jaderberg, D. Hassabis, J. M. Jumper, Accurate structure prediction of biomolecular interactions with AlphaFold 3. Nature 630, 493–500 (2024).

37. D. Listov, S. J. Fleishman, *De novo* enzyme design: Controlling structure to design function. Current Opinion in Structural Biology 98, 103252 (2026).

38. P. A. Romero, F. H. Arnold, Exploring protein fitness landscapes by directed evolution. Nat Rev Mol Cell Biol 10, 866–876 (2009).

39. V. Q. Tran, M. Nemeth, L. J. Bartie, S. S. Chandrasekaran, A. Fanton, H. C. Moon, B. L. Hie, S. Konermann, P. D. Hsu, Rapid directed evolution guided by protein language models and epistatic interactions. Science 0, eaea1820 (2026).

40. S. C. Hammer, A. M. Knight, F. H. Arnold, Design and evolution of enzymes for non-natural chemistry. Current Opinion in Green and Sustainable Chemistry 7, 23–30 (2017).

41. J. Gu, Y. Xu, Y. Nie, Role of distal sites in enzyme engineering. Biotechnology Advances 63, 108094 (2023).

42. N. Zarifi, P. Asthana, H. Doustmohammadi, C. Klaus, J. Sanchez, S. E. Hunt, R. V. Rakotoharisoa, S. Osuna, J. S. Fraser, R. A. Chica, Distal mutations enhance catalysis in designed enzymes by facilitating substrate binding and product release. Nat Commun 16, 8662 (2025).

43. J. Su, C. Han, Y. Zhou, J. Shan, X. Zhou, F. Yuan, SaProt: Protein Language Modeling with Structure-aware Vocabulary. bioRxiv [Preprint] (2023). 10.1101/2023.10.01.560349.

44. C. M. Miton, K. Buda, N. Tokuriki, Epistasis and intramolecular networks in protein evolution. Current Opinion in Structural Biology 69, 160–168 (2021).

45. C. Fröhlich, H. A. Bunzel, K. Buda, A. J. Mulholland, M. W. van der Kamp, P. J. Johnsen, H.-K. S. Leiros, N. Tokuriki, Epistasis arises from shifting the rate-limiting step during enzyme evolution of a β-lactamase. Nat Catal 7, 499–509 (2024).

46. P. G. Mikhael, I. Chinn, R. Barzilay, CLIPZyme: Reaction-Conditioned Virtual Screening of Enzymes. arXiv arXiv:2402.06748 [Preprint] (2024). 10.48550/arXiv.2402.06748.

47. X. Zhang, K. N. Houk, Why Enzymes Are Proficient Catalysts: Beyond the Pauling Paradigm. Acc. Chem. Res. 38, 379–385 (2005).

48. P.-S. Huang, S. E. Boyken, D. Baker, The coming of age of de novo protein design. Nature 537, 320–327 (2016).

49. J. Jiang, G. Yang, F. Ma, Fluorescence coupling strategies in fluorescence-activated droplet sorting (FADS) for ultrahigh-throughput screening of enzymes, metabolites, and antibodies. Biotechnology Advances 66, 108173 (2023).

50. Z. Zou, I. Kalvet, B. Lozhkin, E. Morris, K. Zhang, D. Chen, M. L. Ernst, X. Zhang, D. Baker, T. R. Ward, De novo design and evolution of an artificial metathase for cytoplasmic olefin metathesis. Nat Catal 8, 1208–1219 (2025).

51. M. T. Donato, M. J. Gómez-Lechón, “Fluorescence-Based Screening of Cytochrome P450 Activities in Intact Cells” in Cytochrome P450 Protocols, I. R. Phillips, E. A. Shephard, P. R. Ortiz de Montellano, Eds. (Humana Press, Totowa, NJ, 2013; 10.1007/978-1-62703-321-3_12), pp. 135–148.

52. G. J. Rocklin, T. M. Chidyausiku, I. Goreshnik, A. Ford, S. Houliston, A. Lemak, L. Carter, R. Ravichandran, V. K. Mulligan, A. Chevalier, C. H. Arrowsmith, D. Baker, Global analysis of protein folding using massively parallel design, synthesis, and testing. Science 357, 168–175 (2017).

53. L. Cao, B. Coventry, I. Goreshnik, B. Huang, W. Sheffler, J. S. Park, K. M. Jude, I. Marković, R. U. Kadam, K. H. G. Verschueren, K. Verstraete, S. T. R. Walsh, N. Bennett, A. Phal, A. Yang, L. Kozodoy, M. DeWitt, L. Picton, L. Miller, E.-M. Strauch, N. D. DeBouver, A. Pires, A. K. Bera, S. Halabiya, B. Hammerson, W. Yang, S. Bernard, L. Stewart, I. A. Wilson, H. Ruohola-Baker, J. Schlessinger, S. Lee, S. N. Savvides, K. C. Garcia, D. Baker, Design of protein-binding proteins from the target structure alone. Nature 605, 551–560 (2022).

54. A. Zanghellini, L. Jiang, A. M. Wollacott, G. Cheng, J. Meiler, E. A. Althoff, D. Röthlisberger, D. Baker, New algorithms and an in silico benchmark for computational enzyme design. Protein Science 15, 2785–2794 (2006).

55. F. Richter, A. Leaver-Fay, S. D. Khare, S. Bjelic, D. Baker, De Novo Enzyme Design Using Rosetta3. PLOS ONE 6, e19230 (2011).

56. J. Su, Z. Li, T. Tao, C. Han, Y. He, F. Dai, Q. Yuan, Y. Gao, T. Si, X. Zhang, Y. Zhou, J. Shan, X. Zhou, X. Chang, S. Jiang, D. Ma, M. Steinegger, S. Ovchinnikov, F. Yuan, Democratizing protein language model training, sharing and collaboration. Nat Biotechnol, 1–7 (2025).

57. M. Fey, J. E. Lenssen, FAST GRAPH REPRESENTATION LEARNING WITH PYTORCH GEOMETRIC. (2019).

58. R. J. L. Townshend, M. Vögele, P. Suriana, A. Derry, A. Powers, Y. Laloudakis, S. Balachandar, B. Jing, B. Anderson, S. Eismann, R. Kondor, R. B. Altman, R. O. Dror, ATOM3D: Tasks On Molecules in Three Dimensions. arXiv arXiv:2012.04035 [Preprint] (2022). 10.48550/arXiv.2012.04035.

59. P. G. Mikhael, I. Chinn, R. Barzilay, CLIPZyme: Reaction-Conditioned Virtual Screening of Enzymes. arXiv arXiv:2402.06748 [Preprint] (2024). 10.48550/arXiv.2402.06748.

60. S. Moon, E. Ceyhan, U. A. Gurkan, U. Demirci, Statistical Modeling of Single Target Cell Encapsulation. PLOS ONE 6, e21580 (2011).

61. S. Quan, A. Hiniker, J.-F. Collet, J. C. A. Bardwell, “Isolation of Bacteria Envelope Proteins” in Bacterial Cell Surfaces: Methods and Protocols, A. H. Delcour, Ed. (Humana Press, Totowa, NJ, 2013; 10.1007/978-1-62703-245-2_22), pp. 359–366.

62. J. C. Klein, M. J. Lajoie, J. J. Schwartz, E.-M. Strauch, J. Nelson, D. Baker, J. Shendure, Multiplex pairwise assembly of array-derived DNA oligonucleotides. Nucleic Acids Res 44, e43 (2016).

63. W. Kabsch, XDS. Acta Crystallogr D Biol Crystallogr 66, 125–132 (2010).

64. D. Liebschner, P. V. Afonine, M. L. Baker, G. Bunkóczi, V. B. Chen, T. I. Croll, B. Hintze, L.-W. Hung, S. Jain, A. J. McCoy, N. W. Moriarty, R. D. Oeffner, B. K. Poon, M. G. Prisant, R. J. Read, J. S. Richardson, D. C. Richardson, M. D. Sammito, O. V. Sobolev, D. H. Stockwell, T. C. Terwilliger, A. G. Urzhumtsev, L. L. Videau, C. J. Williams, P. D. Adams, Macromolecular structure determination using X-rays, neutrons and electrons: recent developments in Phenix. Acta Crystallogr D Struct Biol 75, 861–877 (2019).

65. P. Emsley, K. Cowtan, Coot: model-building tools for molecular graphics. Acta Cryst D 60, 2126–2132 (2004).

66. D. Kim, S. M. Woodbury, W. Ahern, D. Tischer, A. Kang, E. Joyce, A. K. Bera, N. Hanikel, S. Salike, R. Krishna, J. Yim, S. J. Pellock, A. Lauko, I. Kalvet, D. Hilvert, D. Baker, Computational design of metallohydrolases. Nature 649, 246–253 (2026).

67. S. Studer, D. A. Hansen, Z. L. Pianowski, P. R. E. Mittl, A. Debon, S. L. Guffy, B. S. Der, B. Kuhlman, D. Hilvert, Evolution of a highly active and enantiospecific metalloenzyme from short peptides. Science 362, 1285–1288 (2018).

68. M. L. Zastrow, A. F. A. Peacock, J. A. Stuckey, V. L. Pecoraro, Hydrolytic catalysis and structural stabilization in a designed metalloprotein. Nature Chem 4, 118–123 (2012).

69. W. J. Song, F. A. Tezcan, A designed supramolecular protein assembly with in vivo enzymatic activity. Science 346, 1525–1528 (2014).

70. C. M. Rufo, Y. S. Moroz, O. V. Moroz, J. Stöhr, T. A. Smith, X. Hu, W. F. DeGrado, I. V. Korendovych, Short peptides self-assemble to produce catalytic amyloids. Nature Chem 6, 303–309 (2014).

71. Y. Wei, M. H. Hecht, Enzyme-like proteins from an unselected library of designed amino acid sequences. Protein Eng Des Sel 17, 67–75 (2004).

72. J. A. Verpoorte, S. Mehta, J. T. Edsall, Esterase Activities of Human Carbonic Anhydrases B and C. Journal of Biological Chemistry 242, 4221–4229 (1967).

73. X. Zhang, K. N. Houk, Why Enzymes Are Proficient Catalysts: Beyond the Pauling Paradigm. Acc. Chem. Res. 38, 379–385 (2005).

74. B. S. Der, D. R. Edwards, B. Kuhlman, Catalysis by a De Novo Zinc-Mediated Protein Interface: Implications for Natural Enzyme Evolution and Rational Enzyme Engineering. Biochemistry 51, 3933–3940 (2012).

75. H. J. Goren, M. Fridkin, The Hydrolysis of p-Nitrophenylacetate in Water. European Journal of Biochemistry 41, 263–272 (1974).

